# *Pandoravirus celtis* illustrates the microevolution processes at work in the giant Pandoraviridae genomes

**DOI:** 10.1101/500207

**Authors:** Matthieu Legendre, Jean-Marie Alempic, Nadège Philippe, Audrey Lartigue, Sandra Jeudy, Olivier Poirot, Ngan Thi Ta, Sébastien Nin, Yohann Couté, Chantal Abergel, Jean-Michel Claverie

## Abstract

With genomes of up to 2.7 Mb propagated in µm-long oblong particles and initially predicted to encode more than 2000 proteins, members of the Pandoraviridae family display the most extreme features of the known viral world. The mere existence of such giant viruses raises fundamental questions about their origin and the processes governing their evolution. A previous analysis of six newly available isolates, independently confirmed by a study including 3 others, established that the Pandoraviridae pan-genome is open, meaning that each new strain exhibits protein-coding genes not previously identified in other family members. With an average increment of about 60 proteins, the gene repertoire shows no sign of reaching a limit and remains largely coding for proteins without recognizable homologs in other viruses or cells (ORFans). To explain these results, we proposed that most new protein-coding genes were created *de novo*, from pre-existing non-coding regions of the G+C rich pandoravirus genomes. The comparison of the gene content of a new isolate, *P. celtis*, closely related (96% identical genome) to the previously described *P. quercus* is now used to test this hypothesis by studying genomic changes in a microevolution range. Our results confirm that the differences between these two similar gene contents mostly consist of protein-coding genes without known homologs (ORFans), with statistical signatures close to that of intergenic regions. These newborn proteins are under slight negative selection, perhaps to maintain stable folds and prevent protein aggregation pending the eventual emergence of fitness-increasing functions. Our study also unraveled several insertion events mediated by a transposase of the hAT family, 3 copies of which are found in *P. celtis* and are presumably active. Members of the Pandoraviridae are presently the first viruses known to encode this type of transposase.

## 1 Introduction

The Pandoraviridae is a proposed family of giant dsDNA viruses - not yet registered by the International Committee on Taxonomy of Viruses (ICTV) - multiplying in various species of Acanthamoeba through a lytic infectious cycle. Their linear genomes, flanked by large terminal repeats, range from 1.9 to 2.7 Mb in size, and are propagated in elongated oblong particles approximately 1.2 µm long and 0.6 µm in diameter (Fig. S1). The prototype strain (and the one with the largest known genome) is *Pandoravirus salinus*, isolated from shallow marine sediments off the coast of central Chile (Philippe et al., 2013). Other members were soon after isolated from worldwide locations. Complete genome sequences have been determined for *P. dulcis* (Melbourne, Australia) (Philippe et al., 2013), P*. inopinatum* (Germany) (Antwerpen et al., 2015), *P. macleodensis* (Australia), *P. neocaledonia* (New Caledonia), and *P. quercus* (France) (Legendre et al., 2018), and three isolates from Brazil (*P. braziliensis*, *P. pampulha*, and *P. massiliensis*) (Aherfi et al., 2018). A standard phylogenetic analysis of the above strains suggested that the Pandoraviridae family consists of two separate clades (Claverie et al., 2018; Fig. 1). The average proportion of identical amino acids between Pandoraviridae orthologs within each clade is above 70% while it is below 55% between members of the A and B clades. Following a stringent reannotation of the predicted protein-coding genes using transcriptomic and proteomic data, our comparative genomic analysis reached the main following conclusions (Legendre et al., 2018):

1. the uniquely large proportion of predicted proteins without homologs outside of the Pandoraviridae (ORFans) is real and not due to bioinformatic errors induced by the above-average G+C content (>60%) of Pandoravirus genomes;
2. the Pandoraviridae pan genome appears “open” (i.e. unbounded);
3. as most of the genes are unique to each strain are ORFans, they were not horizontally acquired from other (known) organisms;
4. they are neither predominantly the result of gene duplications.

**Figure 1.**
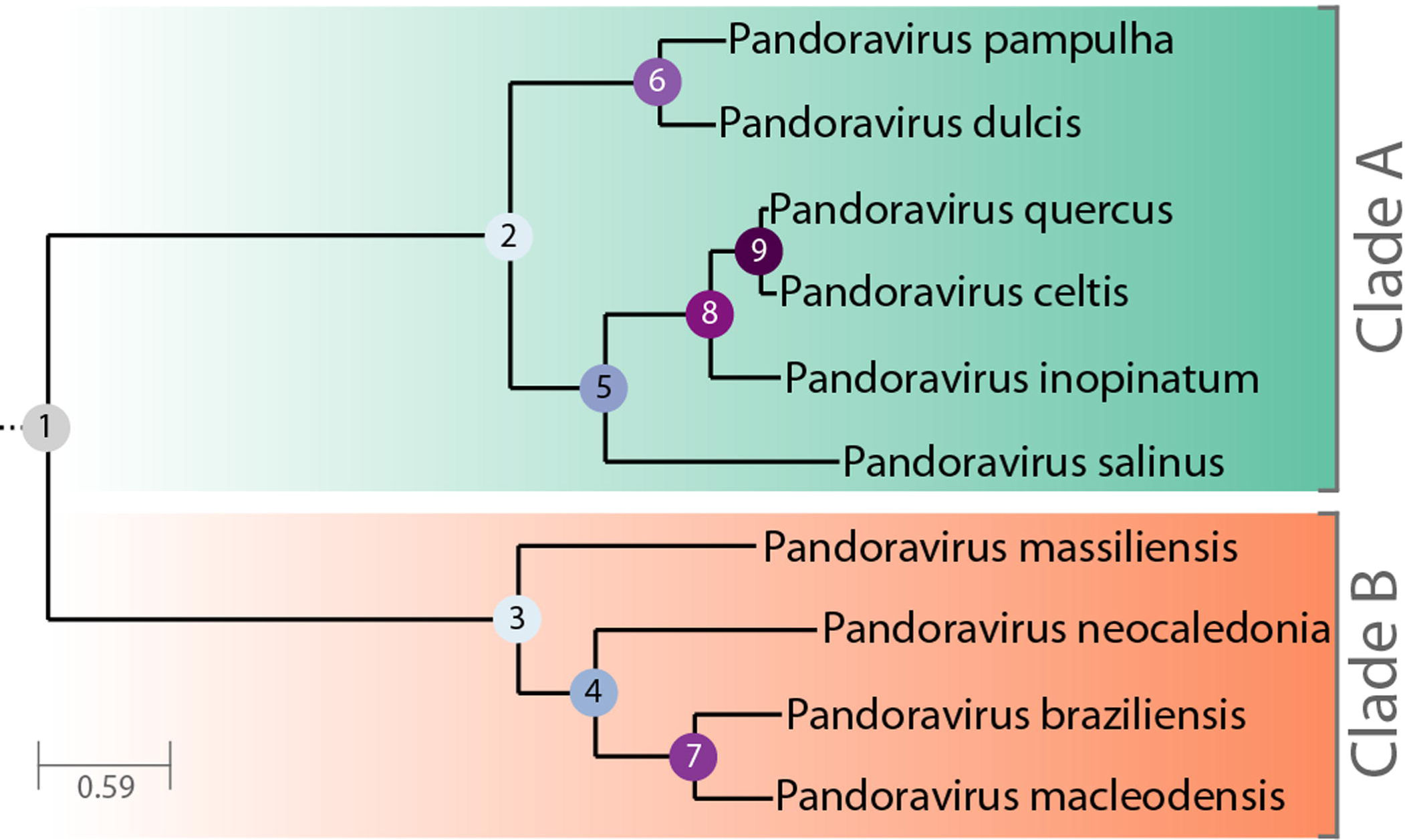
Phylogenetic tree of known, fully characterized Pandoraviruses. The tree was computed from the Pandoraviruses’ core genes (see Materials and Methods for details). The estimated bootstrap values were all equal to 1 and thus not reported. Internal nodes are labeled according to their depth in the tree.

The scenario of *de novo* and *in situ* gene creation, supported by the analysis of their sequence statistical signatures, thus became our preferred explanation for the origin of strain specific genes.

In the present study, we take advantage of the high similarity (96.7% DNA sequence identity) between *P. quercus* and a newly characterized isolate, *P. celtis*, to investigate the microevolution processes initiating the divergence between Pandoraviruses. Our results further support *de novo* gene creation as a main diversifying force of the Pandoraviridae family.

## 2 Materials and Methods

### 2.1 Virus Isolation, Production, and Purification

*P. celtis* and *P. quercus* were isolated in November 2014 from samples of surface soil taken at the base of two trees (*Celtis autralis* and *Quercus ilex*) less than 50 meters apart in an urban green space of Marseille city (GPS: 43°15′16.00″N, 5°25′4.00″E). Their particles were morphologically identical to previously characterized pandoraviruses (Fig. S1). The viral populations were amplified by co-cultivation with *A. castellanii*. They were then cloned, mass-produced and purified as previously described (Philippe et al., 2013).

### 2.2 Genome and transcriptome sequencing, annotation

The *P. celtis* genome was fully assembled from one PacBio SMRT cell sequence data with the HGAP 4 assembler (Chin et al., 2013) from the SMRT link package version 5.0.1.9585 with default parameters and the “aggressive” option=true. Genome polishing was finally performed using the SMRT package. Stringent gene annotation was performed as previously described (Legendre et al., 2018). Briefly, data from proteomic characterization of the purified virions were combined with stranded RNA-seq transcriptomic data, as well as protein homology among previously characterized pandoraviruses. The transcriptomic data were generated from cells collected every hour over an infectious cycle of 15 hours. They were pooled and RNAs were extracted prior poly(A)^+^ enrichment. The RNA were then sent for sequencing. Stranded RNA-seq reads were then used to accurately annotate protein-coding as well as non-coding RNA genes. A threshold of gene expression (median read coverage >5 over the whole transcript) larger than the lowest one associated to proteins detected in proteomic analyses was required to validate all predicted genes (including novel genes) (Table 1). Genomic regions exhibiting similar expression levels but did not encompass predicted proteins or did overlap with protein-coding genes expressed from opposite strands were annotated as “non-coding RNA” (ncRNA) after assembly by Trinity (Grabherr et al., 2011). When only genomic data were available, namely for *P. pampulha*, *P. massiliensi*s and *P. braziliensis* (Aherfi et al., 2018), we annotated protein-coding genes using *ab initio* prediction coupled with sequence conservation information as previously described (Legendre et al., 2018).

**Table 1.**
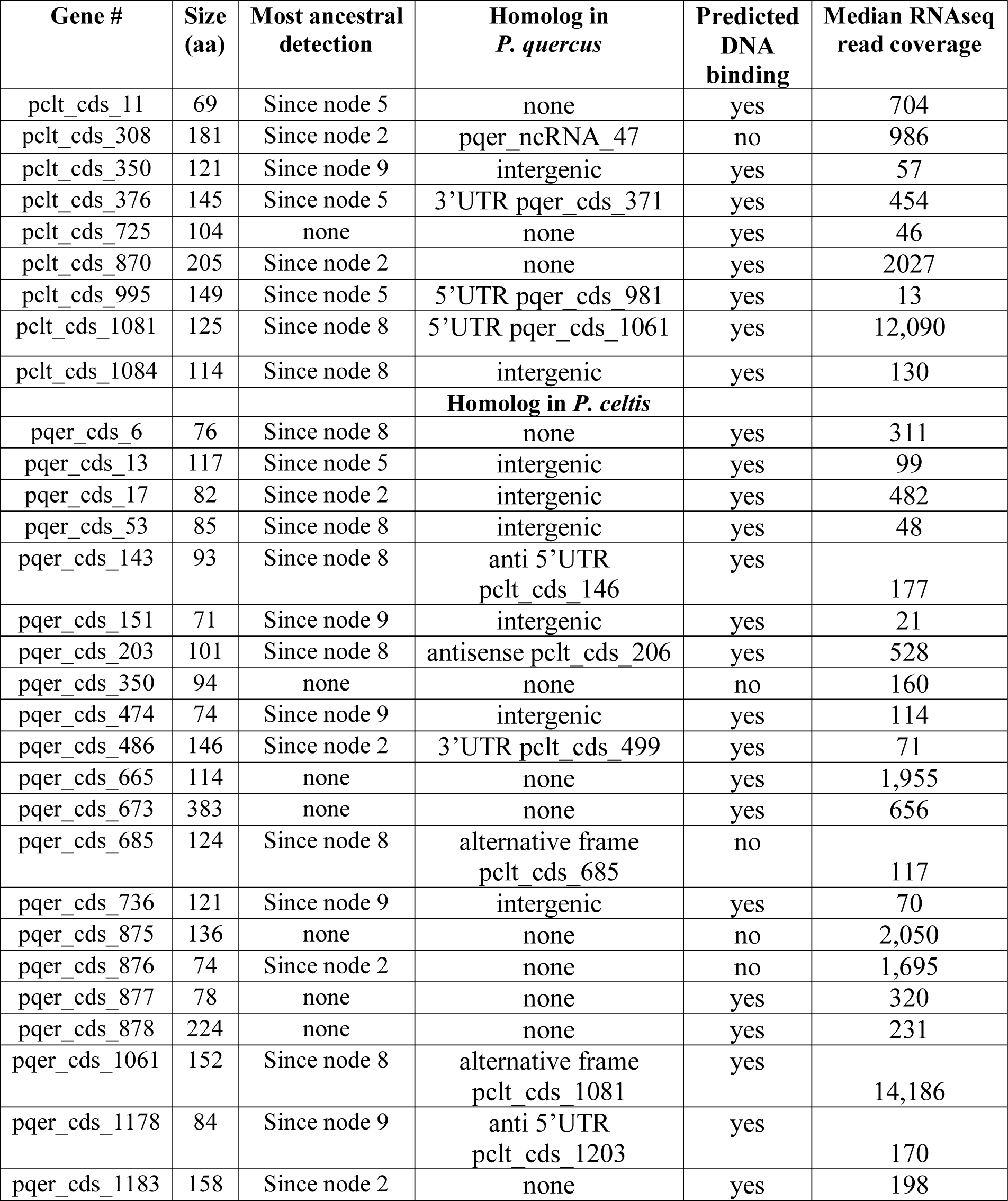
P. celtis and *P. quercus* unique protein-coding genes

Gene clustering was performed on all available Pandoraviruses’ protein-coding genes using Orthofinder (Emms & Kelly, 2015) with defaults parameters except for the “msa” option for the gene tree inference method.

Functional annotation of protein-coding genes was performed using a combination of protein domains search with the CD-search tool (Marchler-Bauer & Bryant, 2004) and HMM-HMM search against the Uniclust30 database with the HHblits tool (Remmert et al., 2011). In addition, we used the same procedure to update the functional gene annotation of *P. salinus*, *P. dulcis*, *P. quercus*, *P. macleodensis* and *P. neocaledonia* (Genbank IDs: KC977571, KC977570, MG011689, MG011691 and MG011690).

### 2.3 Phylogenetic and Selection pressure analysis

The phylogenetic tree (Fig. 1) was computed from the concatenated multiple alignment of the sequences of Pandoravirus core proteins corresponding to single-copy genes. The alignments of orthologous genes peptide sequences were done using Mafft (Katoh et al., 2002). The tree was computed using IQtree (Hoang et al., 2018) with the following options: -m MFP –bb 10000 –st codon –bnni. The best model chosen was: GY+F+R5. Codon sequences were subsequently mapped on these alignments. Ratios of non-synonymous (dN) over synonymous (dS) mutation rates for pairs of orthologous genes were computed using the YN00 method from the PAML package (Yang, 2007). Filters were applied so that dN/dS ratios were only considered if: dN > 0, dS > 0, dS ≤ 2 and dN/dS ≤ 10. We also computed the Codon Adaptation Index (CAI) of *P. celtis* genes using the cai tool from the Emboss package (Rice et al., 2000) as previously described (Legendre et al., 2018).

### 2.4 Particle Proteomics

The *P. celtis* particle proteome was characterized by mass spectrometry-based proteomics from purified viral particles as previously described (Legendre et al., 2018; Legendre et al., 2015).

## 3 Results

### 3.1 Main structural features of the *P. celtis* and *P. quercus* genomes

The *P. celtis* dsDNA genome sequence was assembled as a single 2,028,440 bp-long linear contig, thus slightly shorter than the published 2,077,288-bp for *P. quercus*. Both contains 61% of G+C. The two genomes exhibit a global collinearity well illustrated by a dotplot comparison of their highly similar nucleotide sequences (Fig. 2). In particular, their 48.8 kb difference in genome size does not obviously correspond to a large non-homologous region or a large-scale duplication. Both genomes begin by a nearly perfect 19-kb long palindrome (labelled “P” in Fig. 2). As we did not observe this feature in the other published pandoravirus sequences, it may be specific of *P. celtis* and *P. quercus*, or its absence in other genomes may result from flaws in the assembly of terminal sequences due to insufficient read coverage or quality. Six kb downstream, *P. quercus* exhibits a segment ([25,480-43,420]) nearly identical (20 indels) to the distal end of the genome, inverted (labelled “T” in Fig. 2). Remnants of a similar feature appear blurred in *P. celtis*, but are absent from the other pandoravirus genomes. As these regions are accurately determined, we can infer that a duplication followed by an inversion/translocation of the distal genome terminus occurred in the ancestor of *P. quercus* and *P. celtis* (between node 8 and 9 in Fig. 1).

**Figure 2.**
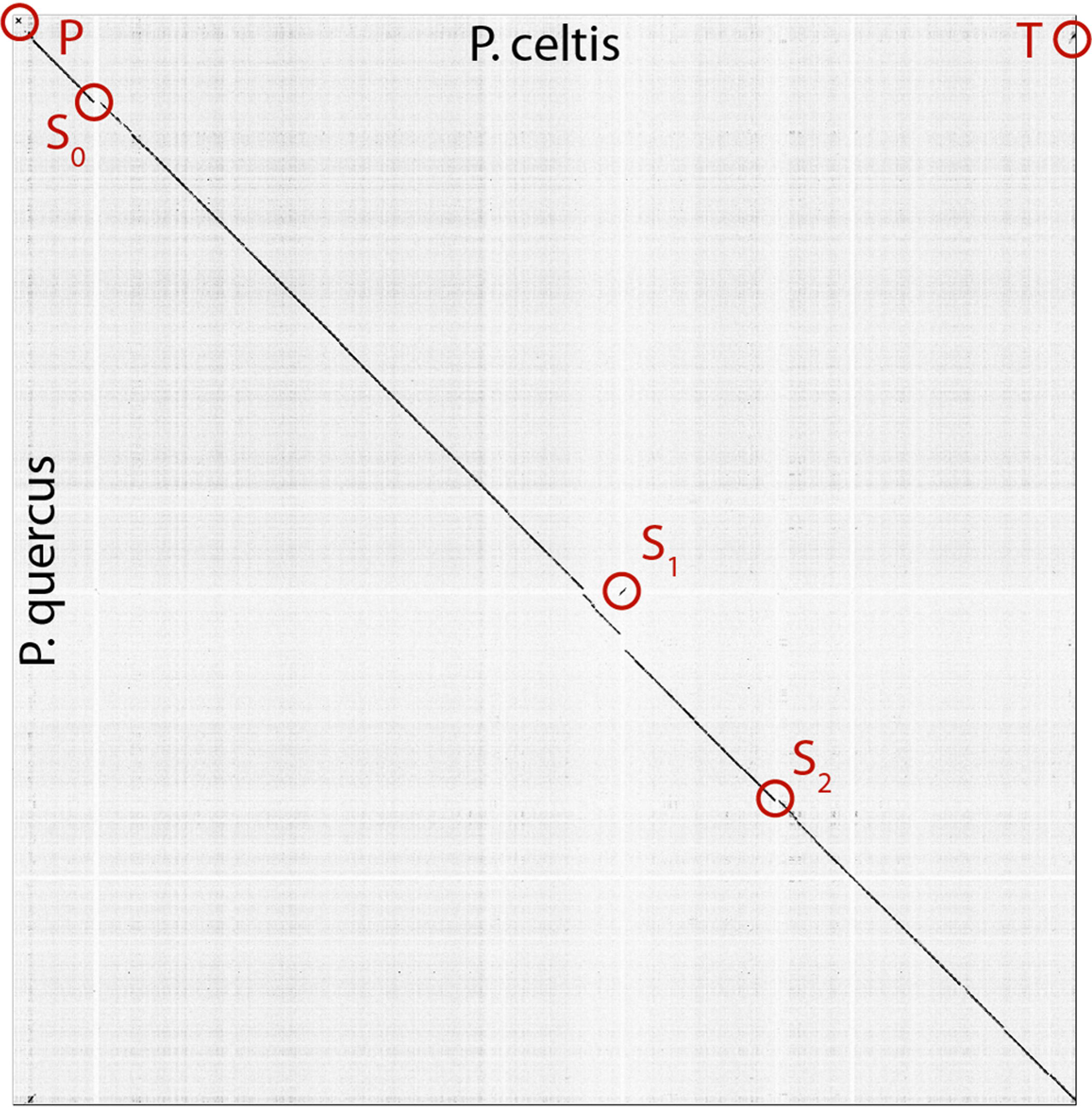
Dot plot (nucleotide) comparison of the *P. celtis vs. P. quercus* genomes. Computations and display were generated using Gepard (Krumsiek et al., 2007). Specific regions, labeled P, T, S_0_, S_1_ and S_2_ are described in the main text.

The next noticeable structural rearrangements consist of three segments denoted S_0_, S_1_, and S_2_ in Fig.2. Each of these segments are flanked by terminal inverted repeats (Fig. S2) and encode a protein (respectively pclt_cds_98, pclt_cds_672, and pclt_cds_871) exhibiting both a BED zinc finger domain and a C-terminal dimerisation region, typical of transposases of the hobo/Activator/Tam3 (hAT) family. All these proteins exhibit the intact signature of hAT transposases and are thus probably active (Atkinson, 2015). Using these sequences as queries, we readily identified other well-conserved hAT transposase homologs in *P. pampulha* (ppam_cds_67, cds_531, cds_663), *P. macleodensis* (pmac_cds_424, cds_799, cds 869) and *P. neocaledonia* (pneo_cds_387, cds_113, cds_658, cds_798).

Besides their common transposase, the S_0_ [10.3 kb, pos. 152,985-163,291], S_1_ [10.3 kb, pos. 1,156,443-1,166,965], and S_2_ [7.3 kb, pos. 1,452,773-1,460,053] transposons encode different sets of proteins. S_0_ encodes 9 proteins (pclt_cds_90-98). Except for the transposase, all of them have no predicted function, and no recognizable homologs outside of the Pandoraviridae (i.e. they are “family ORFans”). S_1_ encodes 11 proteins (pclt_cds_672-682) all of which also have no functional attributes and are family ORFans, except for the transposase. S_2_ encodes 7 proteins (pclt_cds_865-871), all of which have no recognizable signature (except for the transposase and a F-box domain for pclt_cds_870). Besides the transposase, a single protein have paralogs in the S_0_, S_1_, and S_2_ transposons (pclt_cds_92, 681, 869), and one is only shared by S_0_ and S_1_ (pclt_cds_94, 678). These differences clearly suggest that S_0_, S_1_, S_2_ are not the results of recent duplication/transposition events from a common template.

A tentative scenario for the insertion of the *P. celtis* hAT transposons was inferred from their presence/absence in *P. quercus* and the sequence similarity of the transposases. The S_1_ transposon is the only one shared between the two strains (Fig. S2). Moreover, all the orthologous proteins encoded in S_1_ are 100% identical (pclt_cds_672-682 vs. pqer_cds_619-610), including the transposases (pclt_cds_672 and pqer_cds_619). A first possibility is that the S_1_ segment was already present in the ancestor of *P. celtis* and *P. quercus* (Fig. 1, node 9), then was inverted and translocated about 70 kb downstream from its initial location in *P. celtis*. However, since this transposon is absent from *P. inopinatum* (diverging after node 8, Fig. 1), it may have been independently gained from the same source into *P. celtis* and *P. quercus* just after their divergence as two variants within the local viral population. Interestingly, an unrelated sequence of 30.5 kb was inserted at the homologous positions in *P. quercus* (pos. 1,177,379-1,207,935) (Fig. S2). This insertion encodes 24 proteins (pqer_cds_659-682) 13 of which are anonymous and ORFans (i.e. only homologous to other Pandoravirus proteins), the other exhibiting uninformative motifs such as ankyrin repeats (pqer_cds_668, 669, 674-676, 680), F-box domains (pqer_cds_681, 682), Morn repeat (pqer_cds_661) and Ring domain (pqer_cds_670). One protein (pqer_cds_673) exhibits a low (E<10^−2^) and partial similarity with a domain found at the N terminus of structural maintenance of chromosomes (SMC) proteins. However, none of the proteins encoded by this insertion bears any similarity with a hAT family transposase making the mechanism and the origin of this insertion all the more puzzling.

Finally, the proposed scenario concerning S_0_ and S_2_ are simpler. As the two corresponding transposases (pclt_cds_98 and pclt_cds_871) are only 91% identical to each other and less than 67% identical to pclt_cds_672, we propose that they resulted from two independent insertion events that occurred after the *P. quercus*/*P. celtis* divergence, from distinct templates with little overlap in their gene cargo (pclt_cds_92 and 869) besides active transposases. An alternative scenario, would assume the presence of S_0_ and S_2_ in the *P. quercus*/*P. celtis* ancestor, and their subsequent excision in *P. quercus*. This scenario is less likely as *P. inopinatum* (Fig. 1, node 8) does not encode any hAT transposase.

### 3.2 Tandem duplications

Approximately 28 kb downstream from the S_2_ transposon (absent in *P. quercus*), the *P. celtis vs*. *P. quercus* dot plot indicates a cluster of closely interspersed direct repeats (Fig. S3) coding for paralogous proteins that all contain a highly conserved N-terminal fascin-like domain (CD00257). This 80-residue long domain is found in actin-bundling/crosslinking proteins. Many copies of these proteins are found in each of the known pandoraviruses. There are 17 (full-length) copies in *P. quercus*, and 14 in *P. celtis*, the paralogs sharing from 100% to 40% identical residues. Using standard phylogenetic analysis each of the *P. celtis* paralogs are unambiguously clustered with *P. quercus* counterparts in 14 orthologous pairs, indicating that the multiplication of these proteins took place prior to their divergence (Fig. S3). As the 3 additional *P. quercus* paralogs (pqer_cds_130, 131 and 870) with no obvious counterpart in *P. celtis* are not very similar to other *P. quercus* paralogs (max is 72% sequence identity between pqer_cds_870 and pqer_cds_869), it is unlikely that they arise from post-divergence duplications. The most parsimonious scenario is thus that their counterparts have been recently lost by *P. celtis*. In a dot plot, the pqer_cds_130 and pqer_cds_131 genes correspond to a 4 kb deletion at position 224,350 in *P. celtis*. The remaining pqer_cds_870 is located in a dense cluster of 6 contiguous paralogs (pqer_cds_866-871) in *P. quercus*, clearly homologous to a similar cluster of 5 contiguous paralogs (pclt_cds_884-888) *in P. celtis* (Fig. S3). The deletion of a short segment of the *P. celtis* genome is visible at this exact position (1,487,300). Another member of this cluster (pclt_cds_886) just entered a pseudogenization process through the several insertion/deletions breaking its original reading frame (Fig. S3).

### 3.3 Core genes and pan genome update

We previously designed a stringent gene annotation scheme to minimize overpredictions in G+C–rich genomes. In particular, genes predicted to encode proteins without database homolog (i.e. ORFans) were required to overlap with detected sense transcripts to be validated (Legendre et al., 2018). According to this protocol, *P. celtis* encodes at least 1184 protein-coding genes (CDS) compared to 1185 protein-coding genes for *P. quercus*. Both encode a single tRNA(Pro). It is worth noticing that despite a strong reduction compared to the number of proteins initially predicted by standard methods (Philippe et al., 2013), the proportion of ORFans (i.e. w/o homolog outside of the Pandoraviridae family) remains high at 68%. This confirms that the lack of database homologs is not caused by bioinformatics overpredictions (Legendre et al., 2018). *P. celtis* and *P. quercus* shared an average of 96% identical residues, as computed from 822 unambiguous orthologous proteins encoded by single copy genes (i.e. w/o paralogs).

We previously showed that the comparison of the first six available Pandoravirus genomes rapidly converged toward a relatively small set of common genes, corresponding to 455 distinct proteins (clusters) (Legendre et al., 2018). An even smaller estimate (352) was subsequently proposed by another laboratory using a different clustering protocol and strains (Aherfi et al., 2018). Now based on the ten fully sequenced Pandoraviruses and an optimized clustering protocol (see Materials & Methods), the Pandoraviridae core gene set was found to contain 464 genes, close to the asymptotic limit suggested by Fig. 3. Including *P. celtis* in the analysis caused the removal of two proteins from the list. Their homologs in *P. quercus* are pqer_cds_672, and pqer_cds_370, two proteins w/o recognizable functional attribute or homolog outside of the Pandoraviridae. Compared to the average number of distinct proteins encoded by individual Pandoravirus genomes, the proportion of those presumably essential is thus less than half. On the other hand, the predicted gene content of *P. celtis* further increased the Pandoraviridae pangenome. Despite its overall close similarity with *P. quercus*, *P. celtis* remains on a growth curve whereby each new randomly isolated Pandoravirus is predicted to add about 60 distinct proteins to the total number of protein clusters already identified in the family (Fig. 3). The process by which such addition could arise was further investigated by a detailed comparison of the genomes of the most similar relatives, *P. quercus* and *P. celtis*.

**Figure 3.**
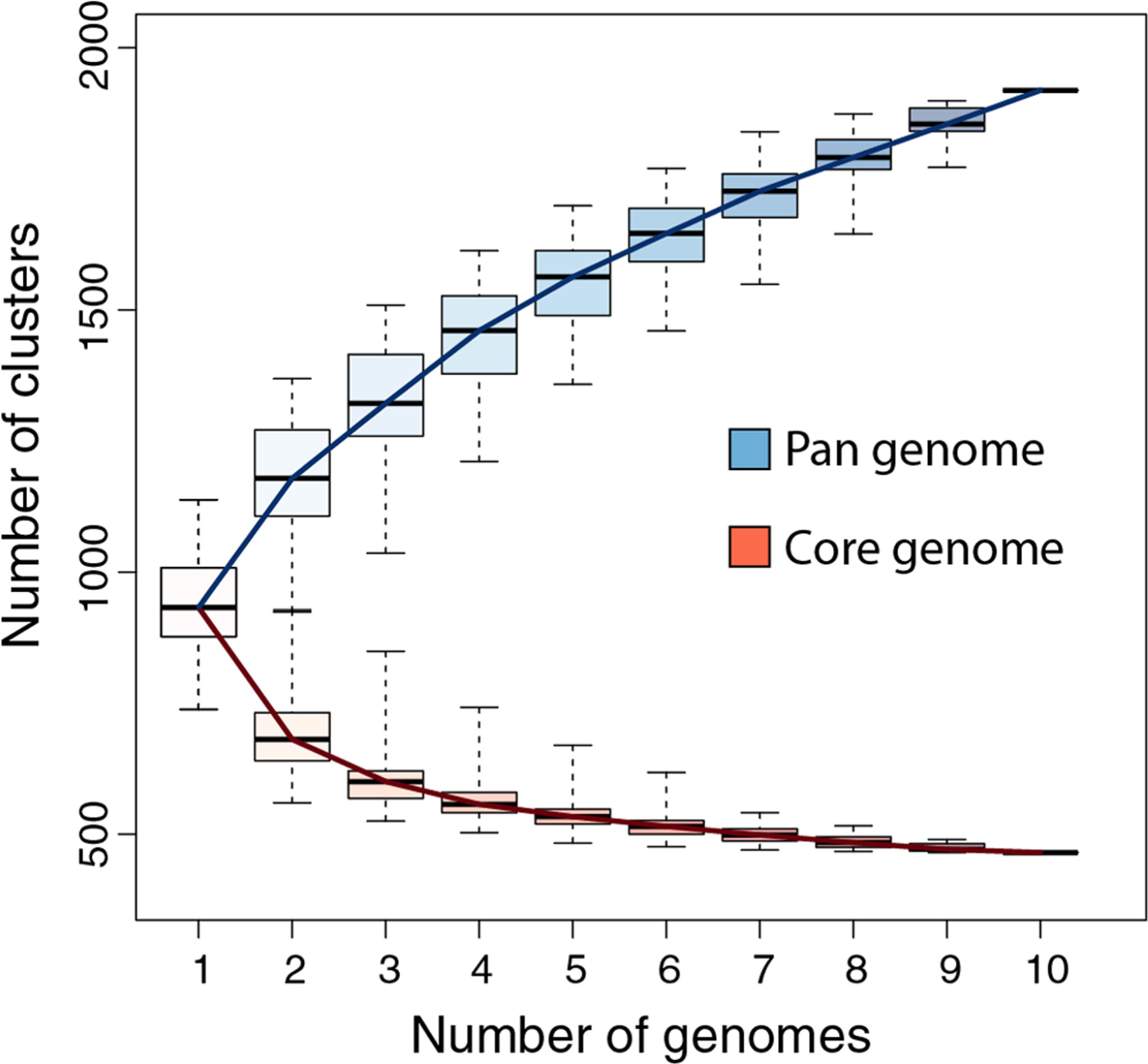
Pandoraviridae pan-genome and core-genome. The boxplot represents the number of protein clusters as a function of the number of sequenced genomes in all possible combinations. The whiskers correspond to the extreme data points.

### 3.4 Protein-coding genes unique to *P. celtis* or *P. quercus*

Given the strong similarity between the two genomes, determining their difference in gene content is straightforward. However, these differences can be due to various mechanisms: the duplication of existing genes, their differential loss, their acquisition by horizontal transfer, or their *de novo* creation. Here we wanted to focus on the later mechanism, previously proposed to be prominent in the Pandoraviridae family (Legendre et al., 2018). The candidate genes most likely resulting from *de novo* creation would be those unique to *P. celtis* or *P. quercus*. If there are no homolog in any of the other Pandoraviridae, *de novo* creation (or acquisition from an unknown source) becomes the most parsimonious evolutionary scenario, compared to a scenario whereby such a gene, present in an ancestral Pandoraviridae would have been independently lost multiple times. The 30 protein-coding genes unique to *P. celtis* (N=9) and *P. quercus* (N=21) are listed in Table 1. As previously noticed (Legendre et al., 2018), these presumed novel genes are associated with statistics that are significantly different from other pandoravirus protein-coding genes. They exhibit a lower Codon Adaptation Index (CAI= 0.233 *vs* 0.351, Wilcoxon test, p < 10^−14^), a lower G+C content (57.5% *vs* 64.4%, p < 10^−16^), and encode smaller proteins (length (aa) =126.6 *vs* 387.3, p < 10^−15^). The two later characteristics make them similar to the random ORFs found in intergenic regions (i.e. the so-called “protogenes”) from which we proposed they originated.

We also noticed that the proteins encoded by the novel genes exhibit an amino-acid composition widely different from the rest of the predicted proteome (Chi-2, df=19, p<10^−10^), with the largest variations observed for lysine (increasing from 2% to 5%), phenylalanine (2.2% to 4%), and arginine (8.6% to 11.3%). Owing to the anomalous proportions of these residues, 25 out of the 30 novel *P. celtis* and *P. quercus* unique proteins are predicted to be DNA-binding (Szilágyi & Skolnick, 2006) (Table 1). Although such predictions are subject to caution (as the predictions remains the same for shuffled sequences) binding novel proteins to the viral genome might help segregate loosely folded (i.e. intrinsically disordered) proteins and diminish their potential interference with the rest of the viral proteome. Over time, these proteins may also gain some regulatory functions.

We investigated the putative intergenic origin of these 30 strain-specific protein-coding genes by searching for remote sequence similarity in the genome of all other Pandoraviruses, using tblastn (protein query against six reading frame translation, E<10^−3^) (Sayers et al., 2018). For seven of them, one unique to *P. celtis* and six unique to *P. quercus*, we found no similarity within the non-coding moiety of other pandoraviruses. Our most parsimonious interpretation is that they represent recent independent additions in each genome by *de novo* creation or transfer from an unknown organism w/o known relative. A slightly less parsimonious scenario would be their addition prior to node 9, followed by differential losses in *P. quercus* or *P. celtis.* Remote but significant (E<10^−3^) similarities were detected for the 25 other novel genes within non-coding regions of other pandoraviruses, all of them members of Clade A (Fig. 1). These positive matches were distributed as followed: five in strains that diverged since node 9, eight in all strains that diverged since node 8, four in all strains that diverged since node 5, and six in all strains that diverged since node 2. We did not detect significant matches in earlier diverging Clade B members. The distribution of these traces in pandoraviruses at various evolutionary distances is interpreted in the discussion section.

### 3.5 ncRNAs

We annotated *P. celtis* 161 ncRNAs (Table S1), a number close to the 157 ncRNAs predicted for *P. quercus* using the same protocol. Such numerous transcripts without protein coding capability were previously noticed for other pandoravirus strains (Legendre et al., 2018). The *P. celtis* ncRNAs vary in length from 234 to 4,456 nucleotides (median= 1384, mean=1413 ± 639). By comparison, the *P. quercus* ncRNAs range from 273 to 5,176 (median=1,189, mean=1299 ± 738). The two distributions are not significantly different (T-test: p > 0.13). The expression levels (in median read coverage) of *P. celtis* ncRNAs vary from 92 to 1608 (median=283) compared to 3665 to 506,994 (median=690) for protein coding transcripts. A similar non-coding *versus* coding expression median ratio (≈ 0.41) was previously found for *P. quercus* (ncRNAs median =228, protein-coding transcript median=551), in a distinct experiment. As previously noticed for other pandoravirus strains, most of *P. celtis* ncRNAs (154/161=95.6%)) are overlapping by more than half of their length with a protein-coding transcript expressed from the opposite strand, and only 7 are mostly intergenic.

Although the *P. celtis* and *P. quercus* genomes share a very large proportion of their protein-coding genes (1146/1184=96.8%), this was not the case for ncRNAs. We found that only 87 of *P. celtis* ncRNAs (87/161=54%) exhibit a homolog among *P. quercus* ncRNAs (Table S1). The 74 *P. celtis* ncRNAs without homologs thus mostly correspond to the lack of detectable transcription in the cognate sequence of the *P. quercus* genome. Unexpectedly, the *P. celtis* and *P. quercus* novel protein-coding genes discussed above (section 3.4) only rarely overlap with ncRNAs in other strains. The single case is pclt_cds_308, the coding region of which overlaps with a *P. quercus* ncRNA (pquer_ncRNA_47) (Table 1).

### 3.6 Selection pressure on new genes

Protein-coding genes shared by at least two different Pandoraviruses provide an opportunity to estimate the selection pressure acting on them by computing the ratio of nonsynonymous (dN) to synonymous (dS) substitutions per site. We computed the dN/dS ratio of genes shared by increasingly close Pandoraviruses, dating their creation or acquisition by reference to the corresponding node in the Pandoravirus phylogenetic tree (nodes 1 to 9 in Fig 1). All pairs of orthologous gene sequences were thus analyzed using the YN00 algorithm from the PAM package (Yang, 2007) and assigned to their most likely creation/acquisition node based on their presence or absence in various clades of Pandoraviruses. As shown in Fig. 4 all genes are on average under negative selection pressure (dN/dS < 1), including the presumably most recently created/acquired genes as those only found in *P. quercus* and *P. celtis* (i.e. node 9 in Fig. 1). As previously documented for short evolutionary distances and very close gene sequences (our case), the computed dN/dS values are probably overestimated (i.e. closer to 1) given that a fraction of the deleterious mutations might not yet be fixed (Rocha et al., 2006). This is consistent with the observed negative correlation between the depth of creation/acquisition and the computed purifying selection (Fig. 4). In other words recently created/acquired genes appear under selective constraints weaker than that of “older” genes. The fact that dN/dS values tend to decrease with longer divergence times was previously noticed (Rocha et al., 2006).

**Figure 4.**
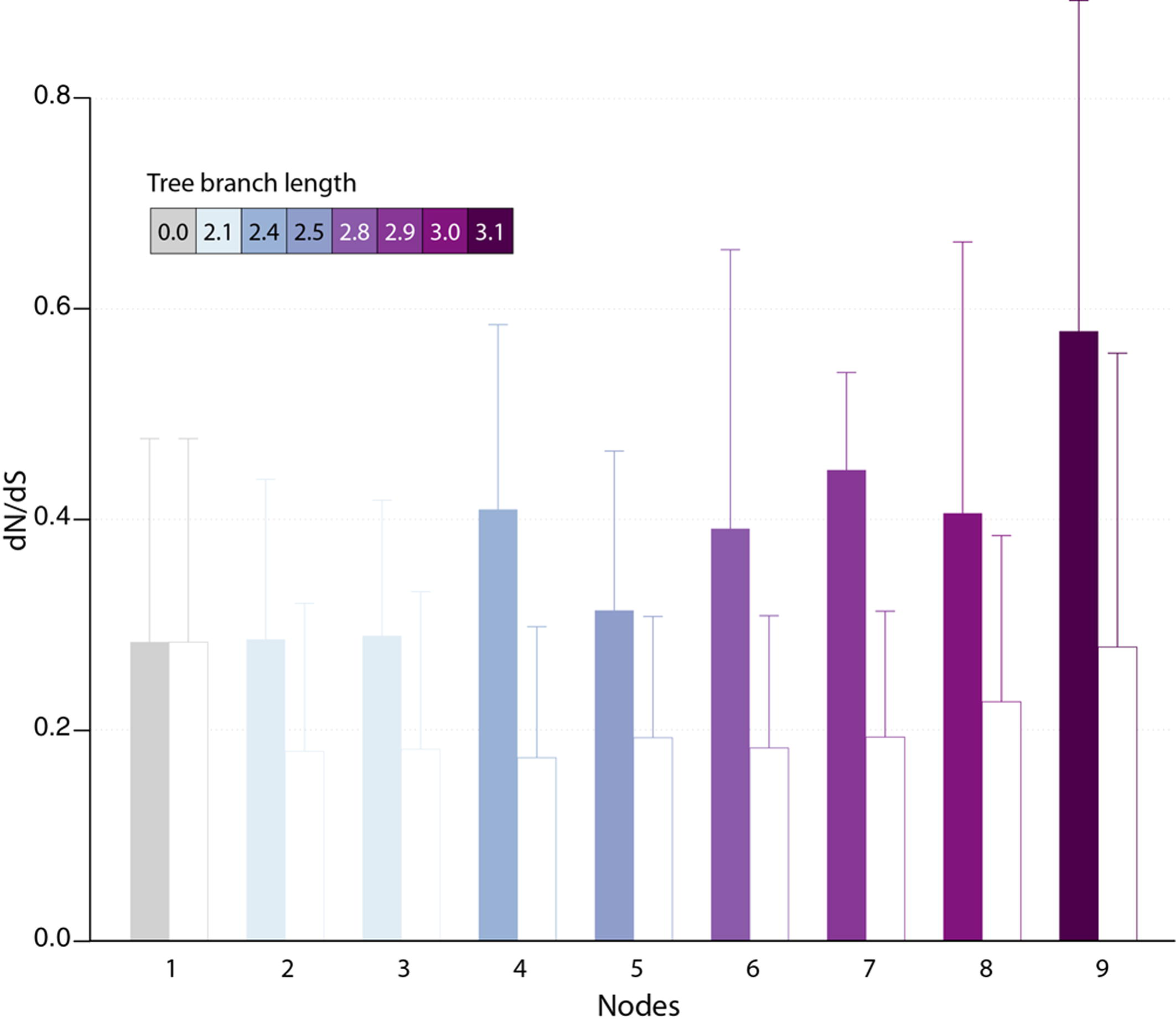
Estimated selection pressure acting on Pandoravirus genes as a function of their ancestry. Shown is the mean of dN/dS ratios (filled bars) for genes that are unique to the pandoravirus strains below a given node in their phylogeny tree (see Fig. 1). Error bars correspond to standard deviations. Bars are colored according to the depth of the node in the tree. As a control, we calculated the dN/dS of Pandoravirus core genes (empty bars). For a given node, we considered pairs of orthologous genes whose common ancestor correspond to that node.

## 4 Discussion

Following our previous analysis of 6 available members of the Pandoraviridae family, we proposed that the gigantism of their genomes as well as the large proportion of ORFans among their encoded proteins was the result of unusual evolutionary mechanisms, including *de novo* gene creation from previously non-coding sequences (Legendre et al., 2018). In this previous study, we compared pandoravirus isolates exhibiting pairwise similarity ranging from 54 to 88%, as computed from a super alignment of their orthologous proteins. At such evolutionary distances, the observed differences are most often the result of multiple and overlapping elementary variation processes the succession of which becomes impossible to retrace. The subsequent isolation of *P. celtis*, a new pandoravirus strain very similar to *P. quercus* (with orthologous proteins sharing 96% of identical residues in average), gave us the opportunity of identifying and estimating the relative contributions of various types of genomic alterations at work during their microevolution from their recent common ancestor.

The global analyses of the gene contents of the 10 members of the Pandoraviridae family available today confirmed previous estimates of the number of core gene clusters (Aherfi et al., 2018; Legendre et al., 2018) at 464. Such a small proportion (less than half) of presumably “essential” genes compared to the total number of proteins encoded by each pandoravirus genome (ranging from 1070 to 1430, using our stringent protocol) (Legendre et al., 2018) raises the question of the origin and utility of so many “accessory” genes. Non-essential genes are normally eliminated from the genomes of obligate intracellular parasites or symbionts through reductive evolution (Floriano et al., 2018; Latorre & Manzano-Marín, 2017; McCutcheon & Moran, 2011; Lopez-Madrigal et al., 2011; Corradi et al., 2010). The contrast is even more puzzling when the size of the Pandoraviridae pan-genome shows no sign of leveling off after reaching 1910 different protein-coding gene clusters, sustaining a trend predicting that each new isolate will contribute 60 additional clusters. Understanding the mechanism by which new genes, – most of which encode ORFan proteins -, appear in the genome of pandoraviruses was the main goal of our study.

As we investigated the most visible alterations of the otherwise nearly perfect collinearity of the *P. celtis* and *P. quercus* genomes, we identified 3 transposons of the hAT family (S_0_, S_1_ and S_2_ in Fig. 2). The cargo of these mobile elements was found to be variable in gene number (10 for S_0_, 11 for S_1_, and 7 for S_2_) and with a single overlap (pclt_cds_92, 681, 869) in addition to the transposases. These by-standing proteins exhibit no functional signature and have no homolog outside of the Pandoraviridae family. Interestingly, hAT family transposases have recently been identified in various Acanthamoeba species (Zhang et al., 2018). However, the gene contents of the *P. celtis* and *P. quercus* hAT transposons indicates that these mobile elements are not prominent vehicles of lateral gene transfers from the amoebal hosts to the pandoraviruses. With the exception of pclt_cds_870 (encoded by S_2_), newly inserted transposons do not contribute genes unique to *P. celtis*. The hAT transposable elements only appear to ferry genes between pandoravirus strains, generating non-local duplications. Such exchanges might occur within an amoebal host undergoing multiple infections. To our knowledge, this is the first identification of hAT family transposons in DNA viruses. The fact that hAT transposons are not present in other well-documented families of large and giant virus infecting Acanthamoeba (Abergel et al., 2015; Aherfi et al., 2016; Fabre et al., 2017; Zhang et al., 2018) suggests that its transfer from host to virus is a rare event, or is specifically linked to Pandoravirus infections.

This work identified 30 genes unique to *P. celtis* or *P. quercus* (Table 1) that we interpreted as encoding novel proteins that appeared after the recent divergence of these two strains from their common ancestor (i.e. below node 9 in Fig. 1). These new genes are uniformly distributed along the *P. celtis* and *P. quercus* genomes, and do not co-localize with large genomic insertions or rearrangements, except for pqer_cds_665 and pqer_cds_673 encoded by the 30.5 kb segments unique to *P. quercus* (section 3.1). As noticed in our previous study (Legendre et al., 2018) all these proteins are strict ORFans (not detected in any other organism, virus or other pandoravirus strains) and exhibit statistical features different from other pandoravirus genes, including a G+C content closer to that of non-coding/intergenic regions. We thus proposed that new proteins could be *de novo* created by the triggering of the transcription and translation of pre-existing non-coding sequences (the so-called “protogenes”). Here we further investigated this hypothesis by looking for eventual similarities of the new protein sequences with non-coding regions in other pandoraviruses at increasing evolutionary distances (Fig. 1). For 8 of the 9 new *P. celtis* proteins and 15 of the 21 new *P. quercus* proteins we detected significant non-coding matches in various pandoravirus strains, in good agreement with their phylogenetic relationships. Trace of 5 new genes were only detected at the level of node 9, 8 at the more ancient node 8, 4 at the node 5, and 6 at the root (node 2) of the clade A pandoraviruses. Although these numbers are small and subject to fluctuations, such a distribution of hits suggests an evolutionary process combining a continuous spontaneous generation of open reading frames, followed by their drift back as non-coding sequences, unless they become transcriptionally active, and fixed as new proteins in a given strain. The maximum value of detected traces at node 8 might thus result from a compromise between the time interval required to generate *de novo* proteins, and the time during which they could retain a detectable similarity with the drifting non-coding regions from which they originated.

According to the above gene creation hypothesis, initially non-coding regions could act as precursors for new proteins, by the random opening of a suitable reading frame, followed by its transcriptional activation. Such a scenario is compatible with the distribution of new gene matches in other pandoravirus strains discussed above, where 8 correspond to intergenic regions, and 5 overlap with UTRs (Table 1). The flexibility of intergenic regions is further attested by the fact that 11 of the new genes have no matches. As shown in Table 1, a single new gene (pclt_cds_308) was found to co-localize with a ncRNA (pqer_ncRNA_47) despite the large number of ncRNAs detected in the *P. celtis* and *P. quercus* genomes. ncRNAs thus do not appear to be necessary intermediates in the emergence of new proteins. The low proportion of conserved ncRNAs between the two close strains (54%) suggests that the on/off expression status of non-coding genomic sequences is fluctuating very fast, and may even be variable at the viral population level.

The fact that a negative selection pressure (dN/dS <1) is associated to the genes appeared since the divergence of *P. celtis* and *P. quercus* decisively reinforces the hypothesis of their *de novo* creation. Firstly, it provides the proof that these genes are truly expressed as *bona fide* proteins, given that differences between synonymous *vs.* non-synonymous mutations cannot be generated within non-translated nucleotide sequences. Any significant deviation from neutrality (dN/dS ≈1) can only be due to a selection process exerted on certain amino acids at certain positions of a true protein in order to improve, preserve, or eliminate its function.

The analysis of proteins whose creation appears most recent (node 9 in Fig. 4) indicates a negative selection pressure whose (probably overestimated) average value (0.58) could correspond to functions slightly increasing the pandoravirus fitness. However, the interpretation of the selection pressure in the context of a function is a problem here, since we previously pointed out that it is highly unlikely that a newly created small protein will have a function (enzymatic, regulatory or structural) in the first place, even less a beneficial one (Legendre et al., 2018).

Thus, we propose an alternative interpretation, based on the knowledge accumulated over the years on the principles governing the stability of proteins. According to our scenario, most new proteins are from the translation of pre-existing non-coding “protogenes” (see 3.4) and the rest from DNA inserts of unknown origin (such as pqer_cds_665 and pqer_cds_673). It is then unlikely that many of these new proteins lacking an evolutionary history will adopt a globular fold (Watters et al. 2006; Monsellier & Chiti, 2007; Tanaka et al., 2010). Most will be toxic by causing nonspecific aggregations within the host-virus proteome and be quickly eliminated through the reversal of these newborn viral genes to an untranscribed/untranslated state. In contrast, proteins whose folded 3-D structures do not prove to be toxic will enter an evolutionary process involving a negative selection pressure promoting the conservation of the amino acids responsible for their stability. Their proportion was estimated at about 34% for 100-200 residue proteins (Miao et al., 2004). In absence of an initial function, the other positions/residues will evolve in a neutral manner. Once “fixed”, the new genes will exhibit an average selection pressure lower than one, combining 1/3 of negative selection with 2/3 of neutrality. Following many studies that have concluded that efficient folding (Watters et al., 2006) and prevention of aggregation are important drivers of protein evolution (see Monsellier & Chiti, 2007), our scenario predicts that many genes specific to pandoraviruses should encode proteins not increasing their fitness, but whose stable 3-D structures may eventually serve as innovative platforms for new functions.

In conclusion, the detailed comparative analysis of *P. celtis* with its very close relative *P. quercus* enabled us to estimate the contributions of various microevolutionary processes to the steady increase of the pan-genome of the proposed Pandoraviridae giant virus family. We first showed that large-scale genomic rearrangements (segmental duplications, translocations) are associated to transposable elements of the hAT family, widespread in metazoans, but until now unique to this family of viruses. However, these mobile elements mostly appear to shuffle pandoravirus genes between strains, without creating new ones or promoting host-to-virus horizontal gene transfers. In contrast with the popular view that horizontal gene transfer plays an important role in the evolution of large DNA viruses (Yutin & Koonin (2013); Koonin et al. 2015; Schulz et al., 2017), this is definitely not the main cause of the inflation of the Pandoraviridae genomes, as we previously argued (Claverie & Abergel, 2013; Philippe et al., 2013; Abergel et al., 2015; Legendre et al., 2018). Finally, we also found that locally repeated regions are the siege of a competition between tandem duplication and gene deletion, locally reshaping the genomes without contributing to net genetic innovation (Fig. S3).

In continuity with our previous work on more distant pandoravirus strains, we found that the 30 protein-coding genes born since the divergence between the very close *P. quercus* and *P. celtis* strains were derived from preexisting non-coding sequences or small DNA segments of unknown origins inserted at randomly interspersed locations. We propose that random ORFs constantly emerge in non-coding regions and that their transcription is turned on in some pandoravirus strains, while they remain silent until they are deleted or diverge beyond recognition in others strains. Our results add strong support to the constant *de novo* creation of proteins, few of which are retained with a little initial impact on the virus fitness until eventually acquiring a selectable function.

Such a scenario, particularly visible and active in the Pandoraviridae, might also apply to the large proportion of ORFans encoded by other DNA viruses, from large eukaryotic viruses (Abergel et al., 2015) to much smaller bacteriophages, in which they can be the target of global functional studies (Berjon-Otero et al., 2017). Interestingly, the *de novo* gene creation scenario is gaining more and more acceptance beyond the realm of virology, recently to explain the origin of orphan protein even in mammals (Schmitz et al., 2018).

## 5 Author Contributions

CA, ML, JMC, and YC designed the experiments. JMA, AL, NP, SJ, and YC contributed to the data and performed the experiments. ML, SN, NTT, OP, CA and JMC analyzed the data. ML, CA, and JMC wrote the manuscript.

## 8 Acknowledgments

We kindly thank Hua-Hao Zhang for providing us with the hAT-transposon sequence he identified in the *Acanthamoeba castellanii* genome.

## 9 Supplementary Material

The Supplementary Material for this article can be found online at: https://www.frontiersin.org/articles/XXX/full#supplementary-material

## 11 Data Availability Statement

*P. celtis* annotated genome is deposited in GenBank under Accession n° MK174290.

---

**Figure S1. TEM image of an ultrathin section of *P. celtis* and *P. quercus* viral particles.** The structures of the particles of the other Pandoraviridae strains do not exhibit any noticeable difference.

**Figure S2. Details of the *P. celtis* hAT transposons.** Details of the S_0_, S_1_, and S_2_ segments highlighted in Fig. 2. The genomic coordinates of the transposons in *P. celtis* (horizontal) and their homologous locations in *P. quercus* (vertical) are indicated. The cognate *P. celtis* transposase gene name and its approximate location is indicated for each transposon. Other genes are not represented. Homologous segments are missing in *P. quercus* for S_0_ and S_2_ (See section 3.1). The sequences of the transposon boundaries are shown above the dot-plots. The TIRs (Terminally Inverted Repeats) are highlighted in black. Conserved TIRs and TSDs (Target Site Duplications) are depicted using arrows.

**Figure S3. Details of two tandem repeat clusters of fascin-domain containing genes.** See section 3.2 for comments. The dot-plot (horizontal: *P. celtis*, vertical *P. quercus*) and the corresponding phylogenetic tree of the genes illustrate the well-ordered dynamics of this highly repeated regions, alternating expansion, deletions and pseudogenization. For instance, we noticed the presence of two alleles of the pclt_886 proteins in our initial sequence data. The minor one (482 residues, used in this tree) is 96.7% identical to its pqer_868 homolog, while the major one results into a smaller protein (363 residues) diverging after the first 125 residues due to the presence of 4 indels in the rest of the gene. The evolutionary history of the fascin-domain containing proteins was inferred using the Neighbor-Joining method as implemented in Mega (Kumar S, et al. (2018) MEGA X: Molecular Evolutionary Genetics Analysis across computing platforms. Mol. Biol. Evol. 35:1547-1549.). The evolutionary distances were computed using the Poisson correction method. The analysis involved 31 amino acid sequences. All positions containing gaps and missing data were eliminated. There were 323 positions in the final dataset.

