## Supplementary Table and Figures for "*Pandoravirus celtis* illustrates the microevolution processes at work in the giant Pandoraviridae genomes"

**Supplementary Table S1.** Predicted non-coding RNAs (ncRNAs)

| *P. celtis* gene # | Length (nt) | *P. quercus* homolog gene # | % identity | % overlap | BlastN E-value | Median RNAseq Read coverage |
|---|---|---|---|---|---|---|
| pclt_ncRNA_138 | 647 | pqer_ncRNA_139 | 100 | 100 | 0 | 457 |
| pclt_ncRNA_31 | 1432 | pqer_ncRNA_42 | 100 | 100 | 0 | 453 |
| pclt_ncRNA_137 | 1750 | pqer_ncRNA_138 | 99.9 | 100 | 0 | 605 |
| pclt_ncRNA_101 | 1041 | pqer_ncRNA_104 | 99.8 | 100 | 0 | 420 |
| pclt_ncRNA_92 | 585 | pqer_ncRNA_95 | 98.1 | 100 | 0 | 567 |
| pclt_ncRNA_149 | 968 | pqer_ncRNA_145 | 97.9 | 100 | 0 | 569 |
| pclt_ncRNA_146 | 1707 | pqer_ncRNA_144 | 97.7 | 100 | 0 | 381 |
| pclt_ncRNA_18 | 1416 | pqer_ncRNA_24 | 96.9 | 100 | 0 | 389 |
| pclt_ncRNA_60 | 1223 | pqer_ncRNA_68 | 96.7 | 100 | 0 | 359 |
| pclt_ncRNA_83 | 1259 | pqer_ncRNA_82 | 95.1 | 100 | 0 | 226 |
| pclt_ncRNA_28 | 1362 | pqer_ncRNA_39 | 99.1 | 99.7 | 0 | 376 |
| pclt_ncRNA_155 | 1997 | pqer_ncRNA_149 | 99.8 | 99.6 | 0 | 733 |
| pclt_ncRNA_144 | 648 | pqer_ncRNA_142 | 96.9 | 99.5 | 0 | 717 |
| pclt_ncRNA_109 | 2468 | pqer_ncRNA_112 | 100 | 99.4 | 0 | 394 |
| pclt_ncRNA_133 | 4456 | pqer_ncRNA_134 | 97.2 | 99.3 | 0 | 354 |
| pclt_ncRNA_150 | 2157 | pqer_ncRNA_146 | 100 | 99.1 | 0 | 395 |
| pclt_ncRNA_100 | 1363 | pqer_ncRNA_103 | 100 | 99.0 | 0 | 995 |
| pclt_ncRNA_65 | 813 | pqer_ncRNA_70 | 100 | 99.0 | 0 | 253 |
| pclt_ncRNA_30 | 735 | pqer_ncRNA_40 | 97 | 98.9 | 0 | 324 |
| pclt_ncRNA_153 | 1504 | pqer_ncRNA_148 | 100 | 98.7 | 0 | 231 |
| pclt_ncRNA_113 | 1576 | pqer_ncRNA_117 | 94 | 98.7 | 0 | 540 |
| pclt_ncRNA_77 | 854 | pqer_ncRNA_80 | 100 | 98.6 | 0 | 475 |
| pclt_ncRNA_16 | 1651 | pqer_ncRNA_21 | 97.4 | 97.8 | 0 | 541 |
| pclt_ncRNA_159 | 1587 | pqer_ncRNA_153 | 97 | 97.8 | 0 | 263 |
| pclt_ncRNA_4 | 1467 | pqer_ncRNA_6 | 98.4 | 97.7 | 0 | 737 |
| pclt_ncRNA_106 | 3617 | pqer_ncRNA_109 | 98.1 | 97.6 | 0 | 336 |
| pclt_ncRNA_5 | 1758 | pqer_ncRNA_7 | 97.9 | 97.6 | 0 | 362 |
| pclt_ncRNA_121 | 2235 | pqer_ncRNA_127 | 97.1 | 96.6 | 0 | 358 |
| pclt_ncRNA_24 | 2177 | pqer_ncRNA_35 | 100 | 95.7 | 0 | 396 |
| pclt_ncRNA_72 | 1652 | pqer_ncRNA_73 | 99.4 | 95.0 | 0 | 283 |
| pclt_ncRNA_34 | 1089 | pqer_ncRNA_43 | 95.2 | 94.9 | 0 | 129 |
| pclt_ncRNA_152 | 2018 | pqer_ncRNA_147 | 100 | 94.9 | 0 | 299 |
| pclt_ncRNA_41 | 1547 | pqer_ncRNA_49 | 99.9 | 94.6 | 0 | 478 |
| pclt_ncRNA_145 | 2158 | pqer_ncRNA_143 | 95 | 94.3 | 0 | 142 |
| pclt_ncRNA_13 | 2246 | pqer_ncRNA_14 | 95.4 | 93.5 | 0 | 151 |
| pclt_ncRNA_9 | 1912 | pqer_ncRNA_11 | 99.8 | 93.4 | 0 | 388 |
| pclt_ncRNA_98 | 1956 | pqer_ncRNA_99 | 98 | 93.3 | 0 | 431 |
| pclt_ncRNA_6 | 1281 | pqer_ncRNA_8 | 99.4 | 92.3 | 0 | 260 |
| pclt_ncRNA_143 | 1580 | pqer_ncRNA_140 | 99.7 | 91.8 | 0 | 517 |
| pclt_ncRNA_78 | 1415 | pqer_ncRNA_81 | 100 | 91.8 | 0 | 268 |

| | | | | | | |
|---|---|---|---|---|---|---|
| pclt_ncRNA_7 | 659 | pqer_ncRNA_9 | 96.3 | 91.0 | 0 | 279 |
| pclt_ncRNA_10 | 961 | pqer_ncRNA_13 | 96.2 | 90.9 | 0 | 304 |
| pclt_ncRNA_90 | 1813 | pqer_ncRNA_89 | 99.9 | 89.0 | 0 | 238 |
| pclt_ncRNA_129 | 1198 | pqer_ncRNA_131 | 100 | 87.2 | 0 | 404 |
| pclt_ncRNA_91 | 921 | pqer_ncRNA_91 | 100 | 86.3 | 0 | 292 |
| pclt_ncRNA_97 | 1253 | pqer_ncRNA_98 | 95.4 | 85.9 | 0 | 116 |
| pclt_ncRNA_156 | 1369 | pqer_ncRNA_150 | 99.1 | 85.8 | 0 | 211 |
| pclt_ncRNA_53 | 873 | pqer_ncRNA_62 | 99.6 | 85.3 | 0 | 524 |
| pclt_ncRNA_125 | 907 | pqer_ncRNA_130 | 95.2 | 84.8 | 0 | 402 |
| pclt_ncRNA_161 | 1748 | pqer_ncRNA_157 | 98.1 | 84.4 | 0 | 262 |
| pclt_ncRNA_89 | 1706 | pqer_ncRNA_88 | 98 | 84.3 | 0 | 588 |
| pclt_ncRNA_116 | 1731 | pqer_ncRNA_121 | 99.7 | 84.3 | 0 | 350 |
| pclt_ncRNA_158 | 1329 | pqer_ncRNA_151 | 100 | 84.0 | 0 | 218 |
| pclt_ncRNA_124 | 1090 | pqer_ncRNA_129 | 95.7 | 84.0 | 0 | 461 |
| pclt_ncRNA_134 | 1155 | pqer_ncRNA_135 | 99.9 | 83.3 | 0 | 791 |
| pclt_ncRNA_43 | 1966 | pqer_ncRNA_50 | 100 | 82.1 | 0 | 1041 |
| pclt_ncRNA_20 | 556 | pqer_ncRNA_28 | 99.1 | 82.0 | 0 | 733 |
| pclt_ncRNA_102 | 1539 | pqer_ncRNA_105 | 99.8 | 81.5 | 0 | 294 |
| pclt_ncRNA_47 | 2129 | pqer_ncRNA_54 | 99.7 | 78.6 | 0 | 219 |
| pclt_ncRNA_123 | 1141 | pqer_ncRNA_128 | 100 | 78.1 | 0 | 512 |
| pclt_ncRNA_114 | 1287 | pqer_ncRNA_117 | 69.8 | 76.6 | $<10^{-119}$ | 117 |
| pclt_ncRNA_59 | 2415 | pqer_ncRNA_67 | 98 | 75.0 | 0 | 337 |
| pclt_ncRNA_104 | 1835 | pqer_ncRNA_107 | 99.9 | 74.7 | 0 | 261 |
| pclt_ncRNA_112 | 1045 | pqer_ncRNA_116 | 71.2 | 73.9 | $<10^{-97}$ | 153 |
| pclt_ncRNA_58 | 779 | pqer_ncRNA_66 | 100 | 71.9 | 0 | 177 |
| pclt_ncRNA_57 | 1335 | pqer_ncRNA_65 | 96.9 | 71.6 | 0 | 770 |
| pclt_ncRNA_54 | 1035 | pqer_ncRNA_64 | 96.1 | 69.3 | 0 | 311 |
| pclt_ncRNA_3 | 1728 | pqer_ncRNA_5 | 98.6 | 69.0 | 0 | 771 |
| pclt_ncRNA_74 | 1930 | pqer_ncRNA_77 | 85.4 | 68.7 | 0 | 278 |
| pclt_ncRNA_94 | 1478 | pqer_ncRNA_96 | 99 | 60.7 | 0 | 655 |
| pclt_ncRNA_15 | 1715 | pqer_ncRNA_20 | 91 | 60.6 | 0 | 376 |
| pclt_ncRNA_25 | 2986 | pqer_ncRNA_37 | 100 | 59.2 | 0 | 142 |
| pclt_ncRNA_17 | 1359 | pqer_ncRNA_23 | 98.7 | 58.6 | 0 | 153 |
| pclt_ncRNA_88 | 744 | pqer_ncRNA_87 | 100 | 58.2 | 0 | 459 |
| pclt_ncRNA_23 | 793 | pqer_ncRNA_34 | 82.9 | 57.4 | $<10^{-142}$ | 263 |
| pclt_ncRNA_45 | 2238 | pqer_ncRNA_52 | 99.9 | 53.4 | 0 | 219 |
| pclt_ncRNA_67 | 1951 | pqer_ncRNA_71 | 100 | 52.4 | 0 | 379 |
| pclt_ncRNA_46 | 1930 | pqer_ncRNA_53 | 100 | 50.9 | 0 | 213 |
| pclt_ncRNA_105 | 2098 | pqer_ncRNA_108 | 98.5 | 50.5 | 0 | 253 |
| pclt_ncRNA_120 | 1630 | pqer_ncRNA_124 | 95.1 | 49.9 | 0 | 304 |
| pclt_ncRNA_14 | 1458 | pqer_ncRNA_15 | 98.3 | 44.8 | 0 | 1608 |
| pclt_ncRNA_70 | 1331 | pqer_ncRNA_129 | 77.4 | 44.3 | $<10^{-126}$ | 119 |
| pclt_ncRNA_103 | 1921 | pqer_ncRNA_106 | 97.2 | 29.7 | 0 | 531 |
| pclt_ncRNA_36 | 2052 | pqer_ncRNA_46 | 100 | 21.1 | 0 | 484 |
| pclt_ncRNA_160 | 1507 | pqer_ncRNA_153 | 74.3 | 20.2 | $<10^{-51}$ | 230 |
| pclt_ncRNA_128 | 1405 | pqer_ncRNA_131 | 85.2 | 18.8 | $<10^{-82}$ | 200 |

| | | | | | | |
|---|---|---|---|---|---|---|
| pclt_ncRNA_115 | 1384 | pqer_ncRNA_118 | 83.9 | 18.8 | $<10^{-73}$ | 276 |
| pclt_ncRNA_95 | 1361 | none | | | | 189 |
| pclt_ncRNA_122 | 1565 | none | | | | 144 |
| pclt_ncRNA_69 | 1623 | none | | | | 139 |
| pclt_ncRNA_68 | 2508 | none | | | | 207 |
| pclt_ncRNA_147 | 1899 | none | | | | 153 |
| pclt_ncRNA_35 | 234 | none | | | | 235 |
| pclt_ncRNA_99 | 643 | none | | | | 376 |
| pclt_ncRNA_85 | 484 | none | | | | 277 |
| pclt_ncRNA_131 | 1933 | none | | | | 247 |
| pclt_ncRNA_162 | 325 | none | | | | 587 |
| pclt_ncRNA_62 | 399 | none | | | | 301 |
| pclt_ncRNA_108 | 808 | none | | | | 136 |
| pclt_ncRNA_55 | 568 | none | | | | 183 |
| pclt_ncRNA_130 | 593 | none | | | | 466 |
| pclt_ncRNA_29 | 3016 | none | | | | 400 |
| pclt_ncRNA_84 | 436 | none | | | | 193 |
| pclt_ncRNA_61 | 725 | none | | | | 167 |
| pclt_ncRNA_42 | 782 | none | | | | 194 |
| pclt_ncRNA_136 | 753 | none | | | | 238 |
| pclt_ncRNA_87 | 367 | none | | | | 166 |
| pclt_ncRNA_132 | 1582 | none | | | | 357 |
| pclt_ncRNA_107 | 1466 | none | | | | 332 |
| pclt_ncRNA_39 | 978 | none | | | | 274 |
| pclt_ncRNA_119 | 1477 | none | | | | 390 |
| pclt_ncRNA_49 | 750 | none | | | | 230 |
| pclt_ncRNA_117 | 1518 | none | | | | 344 |
| pclt_ncRNA_40 | 803 | none | | | | 132 |
| pclt_ncRNA_64 | 732 | none | | | | 235 |
| pclt_ncRNA_26 | 581 | none | | | | 194 |
| pclt_ncRNA_48 | 643 | none | | | | 184 |
| pclt_ncRNA_142 | 644 | none | | | | 222 |
| pclt_ncRNA_63 | 949 | none | | | | 297 |
| pclt_ncRNA_141 | 1593 | none | | | | 674 |
| pclt_ncRNA_96 | 1319 | none | | | | 223 |
| pclt_ncRNA_157 | 954 | none | | | | 206 |
| pclt_ncRNA_86 | 681 | none | | | | 200 |
| pclt_ncRNA_110 | 874 | none | | | | 352 |
| pclt_ncRNA_118 | 989 | none | | | | 328 |
| pclt_ncRNA_38 | 891 | none | | | | 277 |
| pclt_ncRNA_2 | 1683 | none | | | | 271 |
| pclt_ncRNA_11 | 727 | none | | | | 92 |
| pclt_ncRNA_82 | 1363 | none | | | | 332 |
| pclt_ncRNA_66 | 661 | none | | | | 317 |
| pclt_ncRNA_37 | 909 | none | | | | 172 |
| pclt_ncRNA_51 | 1389 | none | | | | 310 |

| | | | | | | |
|---|---|---|---|---|---|---|
| pclt_ncRNA_44 | 915 | none | | | | 343 |
| pclt_ncRNA_139 | 1580 | none | | | | 165 |
| pclt_ncRNA_127 | 1071 | none | | | | 262 |
| pclt_ncRNA_12 | 1786 | none | | | | 222 |
| pclt_ncRNA_75 | 1435 | none | | | | 218 |
| pclt_ncRNA_22 | 903 | none | | | | 189 |
| pclt_ncRNA_71 | 1037 | none | | | | 250 |
| pclt_ncRNA_52 | 1249 | none | | | | 160 |
| pclt_ncRNA_126 | 935 | none | | | | 207 |
| pclt_ncRNA_140 | 1627 | none | | | | 169 |
| pclt_ncRNA_73 | 794 | none | | | | 240 |
| pclt_ncRNA_163 | 1992 | none | | | | 236 |
| pclt_ncRNA_80 | 1113 | none | | | | 574 |
| pclt_ncRNA_135 | 1174 | none | | | | 196 |
| pclt_ncRNA_111 | 1244 | none | | | | 389 |
| pclt_ncRNA_50 | 1550 | none | | | | 172 |
| pclt_ncRNA_154 | 1378 | none | | | | 133 |
| pclt_ncRNA_8 | 1095 | none | | | | 319 |
| pclt_ncRNA_27 | 2220 | none | | | | 816 |
| pclt_ncRNA_19 | 1935 | none | | | | 278 |
| pclt_ncRNA_151 | 1453 | none | | | | 226 |
| pclt_ncRNA_76 | 1863 | none | | | | 130 |
| pclt_ncRNA_56 | 2488 | none | | | | 229 |
| pclt_ncRNA_1 | 1646 | none | | | | 391 |
| pclt_ncRNA_21 | 2922 | none | | | | 100 |
| pclt_ncRNA_33 | 1821 | none | | | | 659 |
| pclt_ncRNA_81 | 2228 | none | | | | 888 |
| pclt_ncRNA_148 | 1849 | none | | | | 355 |
| pclt_ncRNA_79 | 2325 | none | | | | 226 |

The predicted *P. celtis* ncRNA genes were compared to *P. quercus* genome sequence using BlastN. Previously annotated *P. quercus ncRNA* genes overlapping with each best matching genomic positions were defined as homologous (column 3) to the corresponding *P. celtis* ncRNA genes. The transcript size is given in column 2. The overlap size and % identity are given in columns 4 and 5 (if applicable). As all ncRNA homologs originate from two closely related genomes, strong sequence similarities may not imply the conservation of a function (if any). Most of ncRNAs are antisense to a protein-coding gene except for 7 of them (grey-filled rows) that are mostly intergenic. A little more than half (87/161=54%) of the predicted *P. celtis* ncRNA genes exhibit a homolog in *P. quercus*.

### Supplementary Figures S1-S3

#### Figure S1.

**TEM image of an ultrathin section of *P. celtis* and *P. quercus* viral particles.** The structures of the particles of the other Pandoraviridae strains do not exhibit any noticeable difference.

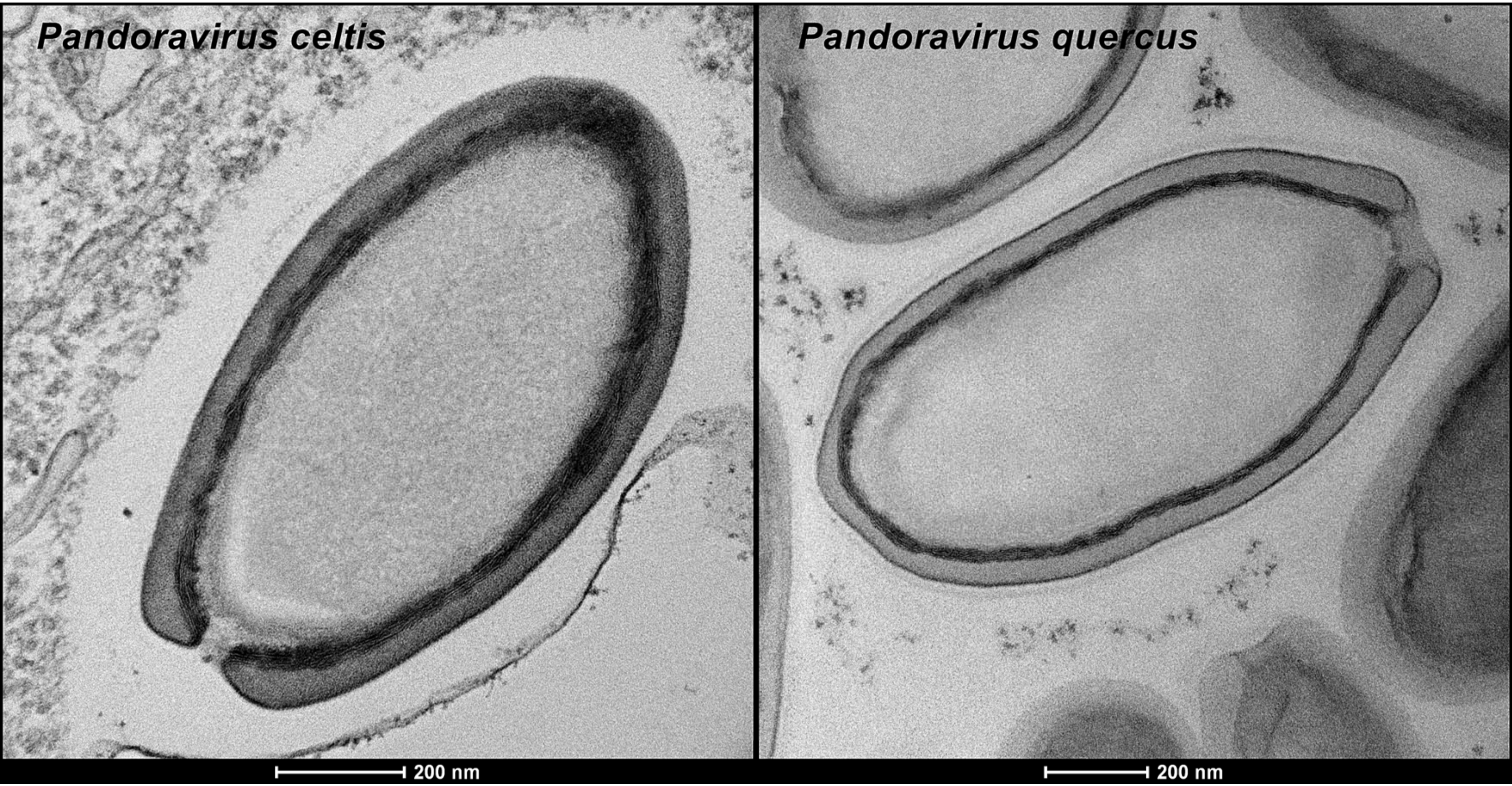
Pandoravirus celtis
Pandoravirus quercus
200 nm
200 nm

#### Figure S2.

**Details of the P. celtis hAT transposons.** Details of the $S_0$, $S_1$, and $S_2$ segments highlighted in Fig. 2. The genomic coordinates of the transposons in *P. celtis* (horizontal) and their homologous locations in *P. quercus* (vertical) are indicated. The cognate *P. celtis* transposase gene name and its approximate location is indicated for each transposon. Other genes are not represented. Homologous segments are missing in *P. quercus* for $S_0$ and $S_2$ (See section 3.1). The sequences of the transposon boundaries are shown above the dot-plots. The TIRs (Terminally Inverted Repeats) are highlighted in black. Conserved TIRs and TSDs (Target Site Duplications) are depicted using arrows.

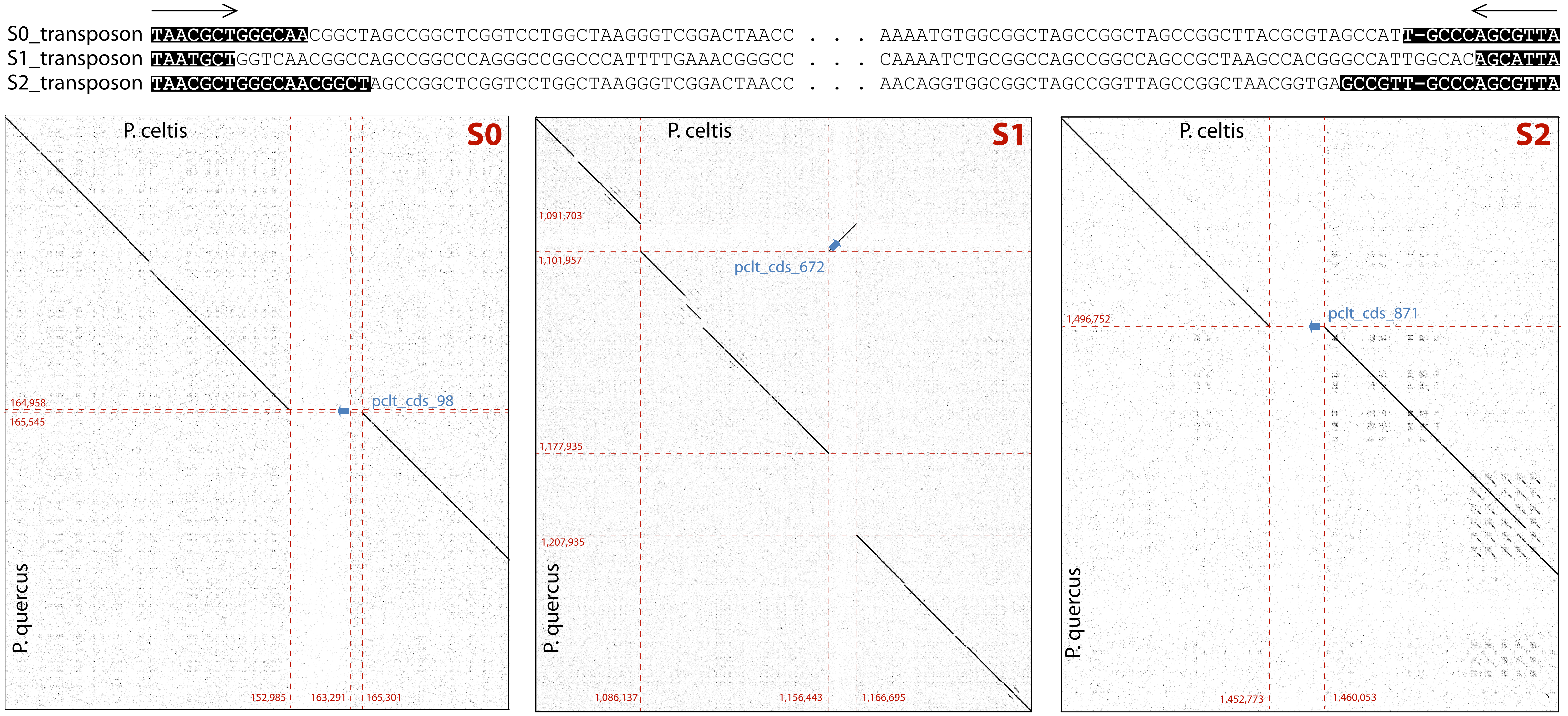
S0_transposon TAACGCTGGGCAACGGCTAGCCGGCTCGGTCCTGGCTAAGGGTCGGACTAACC . . . AAAATGTGGCGGCTAGCCGGCTAGCCGGCTTACGCGTAGCCATT-GCCCAGCGTTA
S1_transposon TAATGCTGGTCAACGGCCAGCCGGCCCAGGGCCGGCCCATTTTGAAACGGGCC . . . CAAAATCTGCGGCCAGCCGGCCAGCCGCTAAGCCACGGGCCATTGGCACAGCATTA
S2_transposon TAACGCTGGGCAACGGCTAGCCGGCTCGGTCCTGGCTAAGGGTCGGACTAACC . . . AACAGGTGGCGGCTAGCCGGTTAGCCGGCTAACGGTGAGCCGTT-GCCCAGCGTTA
P. celtis
S0
P. quercus
164,958
165,545
pclt_cds_98
152,985
163,291
165,301
P. celtis
S1
P. quercus
1,091,703
1,101,957
pclt_cds_672
1,177,935
1,207,935
1,086,137
1,156,443
1,166,695
P. celtis
S2
P. quercus
1,496,752
pclt_cds_871
1,452,773
1,460,053

**Figure S3.**

**Details of two tandem repeat clusters of fascin-domain containing genes.** See section 3.2 for comments. The dot-plot (horizontal: *P. celtis*, vertical *P. quercus*) and the corresponding phylogenetic tree of the genes illustrate the well-ordered dynamics of this highly repeated regions, alternating expansion, deletions and pseudogenization. For instance, we noticed the presence of two alleles of the pclt_886 protein in our initial sequence data. The minor one (482 residues, used in this tree) is 96.7% identical to its pqer_868 homolog, while the major one results into a smaller protein (363 residues) diverging after the first 125 residues due to the presence of 4 indels in the rest of the gene. The evolutionary history of the fascin-domain containing proteins was inferred using the Neighbor-Joining method as implemented in Mega (Kumar S, et al. (2018) MEGA X: Molecular Evolutionary Genetics Analysis across computing platforms. Mol. Biol. Evol. 35:1547-1549.). The evolutionary distances were computed using the Poisson correction method. The analysis involved 31 amino acid sequences. All positions containing gaps and missing data were eliminated. There were 323 positions in the final dataset.

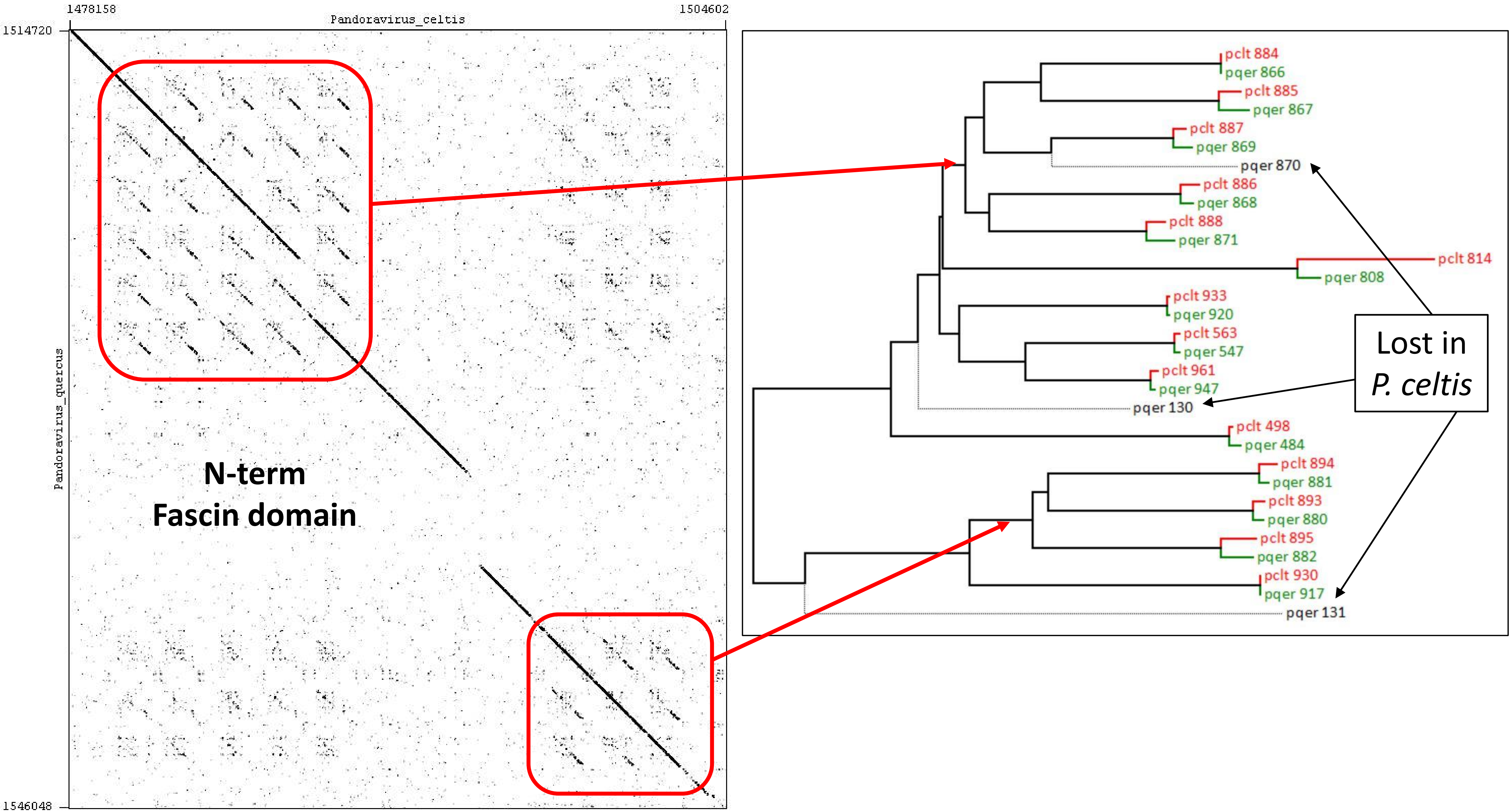
1478158
Pandoravirus_celtis
1504602
1514720
1546048
Pandoravirus_quercus
N-term
Fascin domain
pclt 884
pqer 866
pclt 885
pqer 867
pclt 887
pqer 869
pqer 870
pclt 886
pqer 868
pclt 888
pqer 871
pclt 814
pqer 808
pclt 933
pqer 920
pclt 563
pqer 547
pclt 961
pqer 947
pqer 130
pclt 498
pqer 484
pclt 894
pqer 881
pclt 893
pqer 880
pclt 895
pqer 882
pclt 930
pqer 917
pqer 131
Lost in
P. celtis
